# Genetic and genomic architecture in eight strains of the laboratory opossum *Monodelphis domestica*

**DOI:** 10.1101/2021.09.02.458745

**Authors:** Xiao Xiong, Paul B. Samollow, Wenqi Cao, Richard Metz, Chao Zhang, Ana C. Leandro, John L. VandeBerg, Xu Wang

## Abstract

The gray short-tailed opossum is an established laboratory-bred marsupial model for biomedical research. It serves as a critical species for comparative genomics research, providing the pivotal phylogenetic outgroup for studies of derived vs. ancestral states of genomic/epigenomic characteristics for all eutherian mammal lineages. To characterize the current genetic profile of this laboratory marsupial, we examined 79 individuals from eight established laboratory strains. Double digest restriction-site associated DNA sequencing (ddRAD-seq) and whole-genome resequencing experiments were performed to investigate the genetic architecture in these strains. A total of 66,640 high-quality single nucleotide polymorphisms (SNPs) were identified. We analyzed SNP density, average heterozygosity, nucleotide diversity, and population differentiation parameter Fst within and between the eight strains. Principal component and population structure analysis clearly resolve the strains at the level of their ancestral founder populations, and the genetic architecture of these strains correctly reflects their breeding history. We confirmed the successful establishment of the first inbred laboratory opossum strain LSD (inbreeding coefficient F > 0.99) and a nearly inbred strain FD2M1 (0.98 < F < 0.99), each derived from a different ancestral background. These strains are suitable for various experimental protocols requiring controlled genetic backgrounds and for intercrosses and backcrosses that can generate offspring with informative SNPs for studying a variety of genetic and epigenetic processes. Together with recent advances in reproductive manipulation and CRISPR/Cas9 techniques for *M. domestica*, the existence of distinctive inbred strains will enable genome editing on different genetic backgrounds, greatly expanding the utility of this marsupial model for biomedical research.

## Introduction

The gray short-tailed opossum, *Monodelphis domestica* (also known as the “laboratory opossum”), is the world’s predominant laboratory-bred research marsupial species. In nature, *M. domestica* are widely distributed in southern, central, and western Brazil, eastern Bolivia, and northern Paraguay (Macrini 2004; Carvalho et al. 2011). Adult *M. domestica* typically weigh 60-150g, and males are significantly larger than females. The body length ranges from 70 to 180 mm, and the tail is approximately half the combined head and body length (Costa et al. 2003; Voss and Jansa 2003; Cope et al. 2012). These animals are easily maintained in captivity, breed year-round, and reach sexual maturity relatively rapidly (by six months of age). *M. domestica* serves as a key model for comparative genomics research, providing the pivotal phylogenetic outgroup for studies of derived vs. ancestral states of genomic/epigenomic characteristics for all eutherian mammal lineages.

Marsupials diverged from eutherians ~160 million years ago (Graves and Renfree 2013), and each group exhibits lineage-specific (derived) characteristics that have arisen during their independent evolutionary histories. However, the critical biological functions of major organs, essential genetic and molecular pathways, and fundamental developmental processes are conserved in both lineages. This makes marsupials ideal comparative models for many kinds of research, and they are, therefore, commonly used as “alternative mammals” (Renfree 1981; Samollow 2008) in comparative investigations that span many topics relevant to animal development, physiology, and disease susceptibility.

*M. domestica* is widely recognized as an important model organism for biomedical research (Ley 1987; Saunders et al. 1989; Mikkelsen et al. 2007; Keyte and Smith 2008; Vandeberg and Williams-Blangero 2021), in which it is used for studies in development, physiology, neurobiology, metabolic and infectious disease, immunity, genome structure, function, and evolution. For example, opossum pups are born on embryonic day 14, at the same stage of a 6-week human embryo or E11.5 mouse embryo, which enables many kinds of early embryonic studies that cannot be easily conducted with eutherians, in which access to embryos and fetuses at these stages requires considerable disruption of the gestational environment (Cardoso-Moreira et al. 2019; Mahadevaiah et al. 2020). Marsupials are also valuable for investigating major epigenetic processes such as X-chromosome inactivation (Hornecker et al. 2007; Grant et al. 2012; Rodriguez-Delgado et al. 2014; Wang et al. 2014; Waters et al. 2018; Mahadevaiah et al. 2020) and genomic imprinting (Weidman et al. 2006; Lawton et al. 2008; Das et al. 2012; Douglas et al. 2014; Suzuki et al. 2018). *M. domestica* has been used in many medical and disease studies, such as gene expression during neural development (Dooley et al. 2012; Sears et al. 2012; Pavan et al. 2014; Wheaton et al. 2021), hypercholesterolemia and steatohepatitis (Chan et al. 2010; Chan and VandeBerg 2011; Chan et al. 2012), cancer therapy and prevention (Nair and VandeBerg 2012; Nair et al. 2014), immunogenomics (Parra et al. 2008; Morrissey et al. 2021; Schraven et al. 2021), viral pathogenesis (Thomas et al. 2019), and influence of biological sex on social behavior, individual recognition, and associative learning (Gil et al. 2019).

More than 20 genetic strains of *M. domestica* were developed after the importation of the first wild-caught founders from Brazil and Bolivia in 1978 (VandeBerg and Robinson 1997), and the reference genome of this species was sequenced and assembled in 2007 (Mikkelsen et al. 2007). Although complete pedigree records of all animals produced by the *M. domestica* research colony at the University of Texas, Rio Grande Valley (UTRGV), the oldest and largest *M. domestica* colony in existence and the source colony for all others worldwide, population genetic data for this colony have not been investigated in a systematic, comparative manner. High-resolution sequencing data are required to characterize the genomic and genetic architecture of this colony and to establish a better understanding of the relationships and differences among its individual strains.

Restriction site-associated DNA sequencing (RAD-Seq) (Peterson et al. 2012) provides an effective approach for genetic characterization of these strains. This “reduced-representation” Next-generation sequencing method takes advantage of the sequence specificity of restriction endonucleases, and is an ideal and flexible method for genotyping by extracting a repeatable portion of the genome adjacent to restriction sites, allowing researchers to identify genetic markers across the genome and explore the same subset of genomic regions for many individuals of a species (Baird et al. 2008; Davey and Blaxter 2010; Mastretta-Yanes et al. 2015). Double-digest RAD-seq (ddRAD-seq) was developed based on RAD-seq, with improved robustness and reduced cost (Peterson et al. 2012). ddRAD-seq uses a cocktail of two restriction enzymes for double digestion of DNA, followed by precise size selection that recovers a library consisting of only fragments closely conforming to a desired target size (Peterson et al. 2012). With these advantages, ddRAD-seq has been extensively applied to achieve population-level SNP discovery and high-confidence SNP calling (Liu et al. 2017). It is an efficient method for detecting and describing population structure, hybrid individuals, founder events, biogeographic history, and tagging genomic regions in non-model organisms (Lavretsky et al. 2019).

In the present study, we applied the ddRAD-seq approach to 70 individuals of eight *M. domestica* laboratory strains and discovered 67 thousand informative SNP markers. Together with whole-genome resequencing data from an additional nine opossums, these results provide valuable information on diversity and relatedness between opossum strains that will facilitate further strain development and provide a guide for the efficacious selection of strains for novel uses of this species in future biomedical research applications.

## Materials and Methods

### *Monodelphis domestica* strains and animal selection

Eight laboratory opossum strains were selected for ddRAD-seq experiments: AH11L, ATHHN, ATHL, LSD, LL1, FD2M1, FD2M4, and FD8X. Tissue samples (ear pinna, brain, liver) were collected between 2012 and 2015 from animals maintained at the Texas Biomedical Research Institute under approved IACUC breeding SOP (Table 1 and Table S1) and at Texas A&M University (TAMU) covered by TAMU Animal Use Protocols. For AH11L, ATHHN, ATHL, LSD, LL1, FD2M1, and FD2M4 animals, DNA was extracted from the ear pinna or liver. For FD8X animals, DNA was extracted from the brain or liver. In addition, whole-genome resequencing was performed on gDNA derived from ear pinna of six FD8X and three LSD individuals from the breeding colony at UTRGV between 2016 and 2017 (Table 1 and Table S2).

**Table 1.** The sampling information and breeding history of the eight opossum strains.

| Strain | Number of individuals | Individuals passing QC | F value (inbreeding coefficient) | Heterozygosity range | Heterozygosity mean (SD) | History |
| --- | --- | --- | --- | --- | --- | --- |
| AH11L | 10 | 10 | 0.77-0.80 | (0.00027, 0.00069) | 0.00052 (0.00013) | Partially inbred sibling strains, from 9 founders captured near Exu, Pernambuco, Brazil |
| ATHHN | 10 | 10 | 0.79-0.80 | (0.00026, 0.00051) | 0.00042 (0.00007) |  |
| ATHL | 10 | 10 | 0.93-0.94 | (0.00038, 0.00043) | 0.00036 (0.00006) |  |
| FD2M1 | 10 | 10 | 0.98-0.99 | (0.00040, 0.00059) | 0.00035 (0.00016) | Pop1-Pop2 admixed inbred sibling strains. Pop2 ancestry from 2 founders captured near Piraua, Paraiba, Brazil |
| FD2M4 | 6 | 6 | 0.92-0.93 | (0.00019, 0.00035) | 0.00027 (0.00007) |  |
| FD8X | 12+6* | 6+6* | 0.21-0.23 | (0.00028, 0.00061) | 0.00042 (0.00012) | Pop1-Pop5 admixed strain. Random bred, Pop2 ancestry from 2 founders from Brecha, Bolivia. |
| LL1 | 10 | 8 | 0.40-0.45 | (0.00054, 0.00071) | 0.00060 (0.00005) | Random bred, from the same 9 founders from Exu |
| LSD | 10+3* | 10+3* | >0.99 | (0.00008, 0.00020) | 0.00014 (0.00003) | Inbred strain, from the same 9 founders from Exu |

\*: the number of individuals with whole-genome sequencing data.

### Genomic DNA extraction

DNA for ddRAD-seq was extracted from tissue samples (8 brain, 8 liver, and 62 ear pinnae from the eight laboratory opossum strains) using Qiagen DNA Blood and Tissue Mini kit on a QIAcube automated nucleic acid extraction system following manufacture’s protocol (Qiagen, MD). The nine whole-genome resequencing samples were extracted from ear pinna by mincing the tissue and incubating it overnight in 200 ug/ml Proteinase K at 55 °C with gentle shaking. The solution was RNase treated for 1 hour at 37 °C at a final concentration of 20 ug/ml prior to phase separation by phenol/chloroform (15 minutes each: one phenol, one phenol/chloroform, and one chloroform). pH was adjusted by the addition of NaAc at 1/10 the volume. DNA was precipitated with ice-cold 100% EtOH with a glycogen carrier (0.04 mg per ml of EtOH). The precipitate was transferred into a clean tube and washed with 70% EtOH, briefly dried, and resuspended in 10 mM Tris. The yield and quality of extracted gDNA were checked with NanoDrop and Qubit 3.0 Fluorometer (Thermo Fisher Scientific, USA).

### ddRAD-seq library preparation and sequencing

Prior to library preparation, samples were cleaned, quantified, and normalized as described in (Ballare et al. 2019). Samples were prepared by the double-digest restriction-site associated DNA sequencing method originally described by Poland *et al.* (Poland et al. 2012) with modifications (Yang et al. 2020) using the following specific parameters. A preliminary check of several commonly used restriction enzymes determined that the combination of PstI and NlaIII produced an even distribution of DNA fragments between 300 and 500 bp. Following digestion with PstI and NlaIII, samples were ligated to PstI-compatible P5 adapters and one of 48 unique i5 indexes and NlaIII compatible P7 adapters. The rest of the library preparation proceeded as described in (Yang et al. 2020). Final library pools were assessed for size on a fragment analyzer (Agilent) and quantified by qPCR (Kapa Biosystems, Inc., MA), pooled, and quantified. The prepared DNA libraries were sequenced on an Illumina HiSeq2500 sequencer to generate 125 bp paired-end reads at the Texas A&M AgriLife Research: Genomics and Bioinformatics Service (TxGen, College Station, TX).

### Whole-genome resequencing library preparation and sequencing

Whole-genome resequencing libraries were constructed following the Illumina paired-end DNA library preparation protocol (300-350 bp insert size) for six FD8X individuals and three LSD individuals with Illumina TruSeq DNA library kit. After quality control procedures, the libraries were sequenced on an Illumina HiSeq2500 sequencer.

### Data processing and sequencing read alignment

FastQC (Andrews 2010) was used to assess the quality of raw sequencing data (Table S1-S2). Reads were trimmed using Trimmomatic-0.36 (Bolger et al. 2014) with the following parameters: ILLUMINACLIP Nextera_adapters. fa:2:30:10, LEADING:3, TRAILING:3, SLIDINGWINDOW:4:15, and MINLEN:36. The high-quality filtered reads were aligned to the *M. domestica* reference genome monDom5 (Mikkelsen et al. 2007) using BWA (Li and Durbin 2009) with default settings. Reads mapped to multiple regions in the genome were removed. After removing low-quality bases and sequencing adapter contaminations, an average of 6.33 million reads (98.6% of the total reads) per individual were retained. The average percentage of uniquely mapped reads was 90.8% (Table S1). Eight ddRAD-seq individuals were excluded from the analysis due to the limited amount of retained reads or lower overlap genome mapping rate (Table S1). For the 9 whole-genome resequencing samples, after quality control and read alignment, 127.4 million reads per individual passed quality control, and the percentage of uniquely mapped reads was >92% (Table S2).

### SNP identification and SNP calling in ddRAD-seq and whole-genome resequencing data

*De novo* SNP calling was performed using the BAM files generated from genome alignments with UnifiedGenotyper in the Genome Analysis Toolkit (GATK) (McKenna et al. 2010; DePristo et al. 2011). The variants were filtered using VCFtools (Danecek et al. 2011) according to the thresholds: minimum coverage depth of 6X and minimum alignment quality score of 200. A total of 66,640 high-quality SNPs were identified in the 70 ddRAD-seq datasets retained for analysis (out of 78 individuals sequenced) (see Data S1). Proportions of homozygous and heterozygous SNP positions were computed based on the total number of covered bases, which was defined as covered positions with a minimum sequencing depth of 6X.

### Cross validation of ddRAD-seq SNP calls in *M. domestica* RNA-seq datasets

To validate the quality of the SNP calls from our ddRAD-seq data and to evaluate the reliability of our pipeline, we utilized an independent RNA-seq dataset generated in the reciprocal F1 crosses of strains LL1 and LL2 (Wang et al. 2014). LL2 is a random-bred strain derived from an admixture of founders from Population 1 (Pop1) and Population 2 (Pop2) (Figure 1). Approximately 1.5 billion of 51 bp single-end Illumina sequencing reads were generated from a total of 16 F1 individuals (accession number GSE45211). These RNA-seq reads were aligned to the *M. domestica* reference genome assembly (monDom5) using TopHat v1.4.1 (Trapnell et al. 2009), and SNP calling was performed using SAMtools software (Li et al. 2009) to identify 68,000 SNPs (≥8 X coverage in the RNA-seq data) in the transcripts. The RNA-seq SNP set was compared to the ddRAD-seq calls, and we identified 2,151 SNP positions in both datasets. All SNPs have the same alternative allele except for one at chr1:501281170 (Data S2). Close examination of this SNP position discovered that it is a problematic SNP with a third allele in the RNA-seq data. Therefore, the ddRAD-seq calls have 100% agreement with RNA-seq SNP genotypes, indicating the high accuracy of our ddRAD-seq SNP calls.

**Figure 1.**
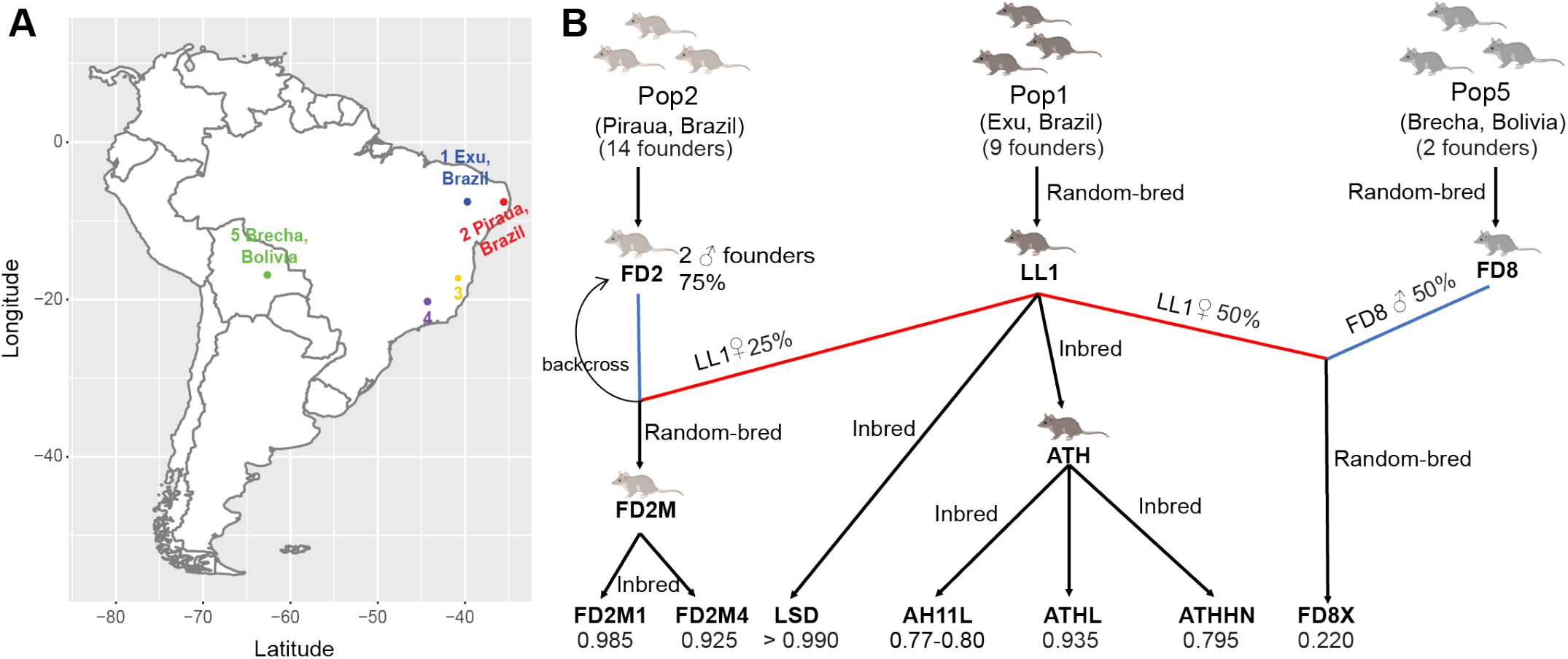
Geographical locations and breeding history of the eight opossum strains in this study. **(A)** Map of South America showing the geographical distribution of five populations. Animals derived from Population 1 (Exu, Brazil), Population 2 (Piraua, Brazil), and Population 5 (Brecha, Bolivia) were sampled for this study. **(B)** Breeding history of eight opossum strains included in this study. Five of the strains were derived from the 9 founders of Population 1. They include a random bred strain, LL1;three partially inbred sibling strains, AH11L, ATHHN, and ATHL: and a fully inbred strain, LSD. Population 2 had 14 founders. Two male Pop2 founders were crossed with LL1 females, followed by backcrossing of F1 females to these two founders, in order to get 75% genetic contribution from Pop2. Subsequent inbreeding resulted in two admixed strains FD2M1 and FD2M4, of which FD2M1is now fully inbred. FD8X is an admixed strain derived from Population 1 and Population 5. The paternal lineage is derived from the sons of one founder male and one founder female captured near the town of Brecha in Bolivia. The maternal lineage is from the LL1 strain. Inbreeding coefficients are indicated at the bottom of this panel.

### Principal component analysis

To examine genetic structure among the eight strains, the SNPs were imported in PLINK (Purcell et al. 2007), and principal component analyses (PCA) were performed using the 70 ddRAD-seq datasets from all eight strains with and without the nine whole-genome resequencing datasets from additional FD8X and LSD animals included. Individual variations in principal components for 66,640 SNP loci of the 70-member ddRAD-seq dataset and the 79-member ddRAD-seq plus nine whole-genomic resequencing dataset were visualized in R.

### Genetic structure analysis

Maximum likelihood estimates of population assignments for each individual were determined using Admixture v1.3 (Alexander et al. 2009; Alexander and Lange 2011; Alexander et al. 2015). Admixture proportions were also estimated using NGSadmix in ANGSD (Allentoft et al. 2015). We performed population structure analyses at *K* = 2, 3, and 4, where *K*= the number of allowed subpopulations. To investigate detailed genetic structure and determine phylogenetic relationships among the strains, we inferred population structure through shared co-ancestry using a model-based Bayesian clustering approach implemented in the fineRADstructure program (Malinsky et al. 2018). A co-ancestry matrix was generated with the RADpainter module in fineRADstructure with default parameters. 100,000 Markov chain Monte Carlo (MCMC) iterations with a burn-in of 100,000 iterations were performed, and sampling occurred every 1,000 iterations to generate the tree file. Finally, a phylogenetic tree of the 70 ddRAD-seq individuals from the eight opossum strains was constructed and visualized in R.

### Estimation of population genetic parameters

The SNP density patterns across each chromosome were plotted using CMplot package (Yin 2018) in R. The Pearson correlation coefficient between inbreeding coefficients (Table S3) and frequencies of heterozygous SNPs in inbred strains were calculated with the Hmisc package (Harrell Jr and Dupont 2006) in R. Composite pairwise estimates of nucleotide diversity (Pi) and genetic variation (Fst Statistic) for autosomal and X-linked ddRAD-seq loci were calculated using the population genomic analysis PopGenome package (Pfeifer et al. 2014) in R with a concatenated data set for 70 ddRAD-seq individuals (Table S3). Results were visualized in 20 kb consecutive windows.

## Results

### Origins and relatedness of laboratory opossum strains examined

Founder animals of the UTRGV colony were imported between 1978 and 1993 from five geographical locations (Figure 1A) in eastern Brazil (Populations 1-4; Pops 1-4) and Bolivia (Population 5, Pop5). Five strains in this study, LL1, LSD, AH11L, ATHHN, and ATHL, were derived from nine Pop1 founders. Among them, AH11L, ATHHN, and ATHL are closely related sibling strains that are partially inbred (Figure 1B). LL1 is a random-bred strain derived from the same nine founders. LSD, which is also derived from the same nine founders, is an inbred strain, which is defined as an animal stock with an inbreeding coefficient (F) in excess of 0.99 (Figure 1B). The FD2M strain was generated by breeding purebred descendants of the two Pop2 founders with LL1 individuals to produce a lineage with 75% Pop2 and 25% LL1 genetic ancestry. FD2M1 and FD2M4 were derived from the inbreeding of FD2M sublines. FD2M1 had a higher inbreeding coefficient (F = 0.985) than FD2M4 (F = 0.925) (Figure 1B and Table 1). FD8X is an admixed strain with maternal genetic background from LL1 animals and paternal background from a Pop5 founder captured near the town of Brecha in Bolivia (Figure 1B). Its calculated ancestry contributions are 50% Pop1:50% Pop5.

### SNP discovery in ddRAD sequencing data

We generated a total of 454 million reads from the 78 samples examined. The ddRAD-seq reads were aligned to the *M. domestica* reference genome monDom5. A total of 70 individuals with sufficient reads and >85% mapping percentage were included in the analysis (Table S1 and S2). In total, we identified 66,640 high-quality SNP positions (Data S1 and S2). The number of SNPs detected per individual varied from 2,817 to 44,739, with an average of 25,013 SNPs per individual. The total length of genomic regions covered by at least 6X ddRAD-seq reads was 54.6 Mbp, indicating a 70-fold genome enrichment. The average genome-wide SNP density was 1.2 SNPs per kb, which is defined as the total number of homozygous and heterozygous SNPs divided by the covered position (depth >= 6). LL1 and the three Pop1-derived partially inbred sibling strains (AH11L, ATHHN, and ATHL) had the lowest proportion of homozygous SNPs (homozygous for the non-reference allele, not including homozygosity for the reference allele). This outcome is in accordance with the fact that the animal sequenced for the *M. domestica* reference genome (Mikkelsen et al. 2007) was from the ATHL strain and had an inbreeding coefficient of 0.917 (VandeBerg, unpublished data) (Figure 2A and Figure S1). The two Pop1/Pop2-admixed strains (FD2M1 and FD2M4) had the highest overall SNP density when compared to the reference genome, which is consistent with their mixed genetic background.

**Figure 2.**
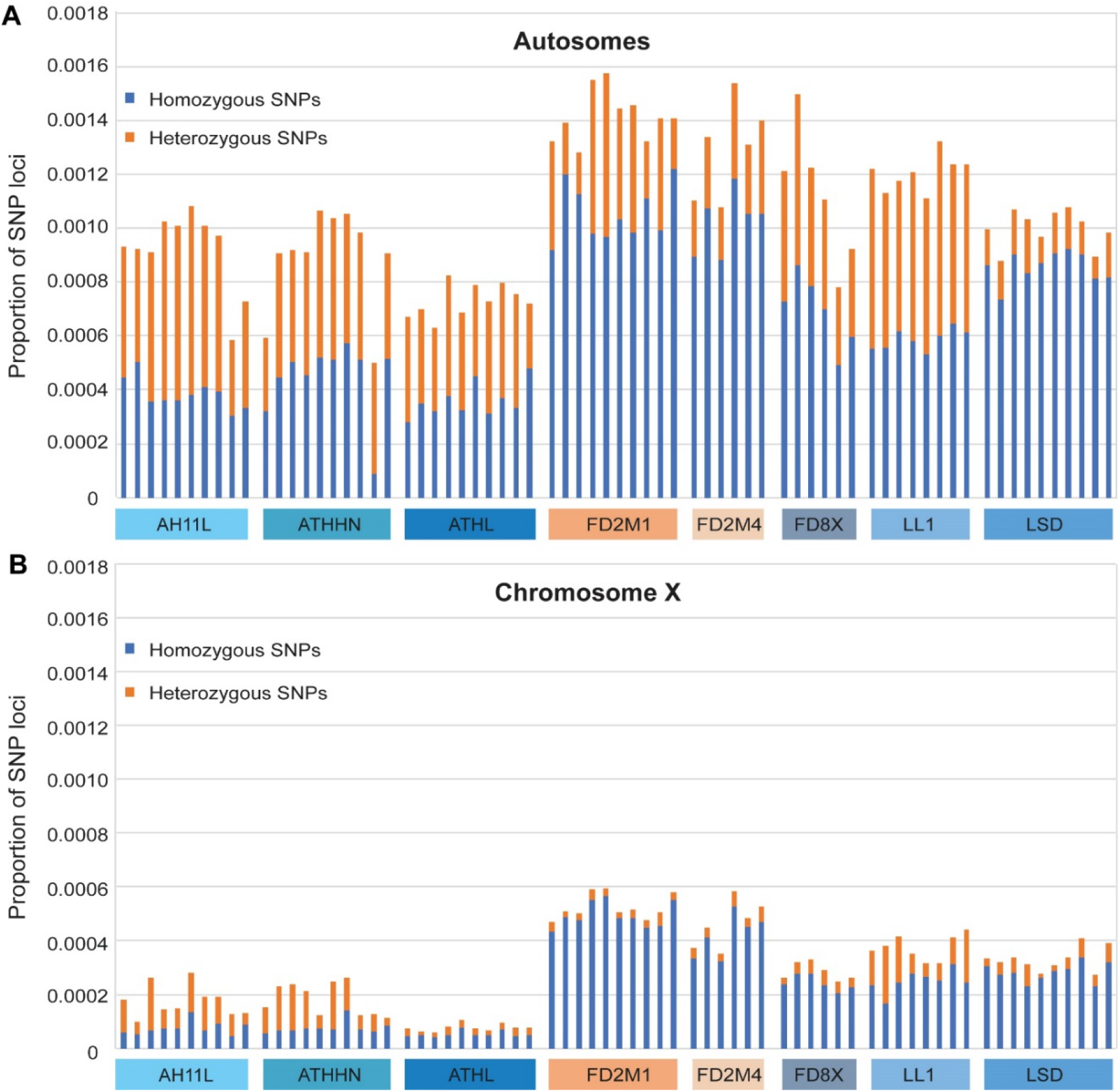
Histograms of proportions of homozygous and heterozygous SNPs for autosomes and the X chromosome of 70 individuals of eight laboratory opossum strains. **(A)** The proportions of autosomal SNPs. Blue: homozygous SNPs; orange: heterozygous SNPs. **(B)** The proportion of X-linked SNPs.

The proportions of both the homozygous and heterozygous X-linked SNPs were significantly less compared to autosomes (Figure 2B). AH11L, ATHHN, and ATHL had lower heterozygosity on the X chromosome compared to the other five strains (Figure 2B). Compared to females, males had a deficiency of heterozygous X-linked SNP loci (Figure S2), which was expected because males are hemizygous. The residual X-linked heterozygous SNPs in males could be due to misassembled autosomal contigs on the X chromosome, multiple copies on the X, or homology between X and autosomal sequences.

### Assessment of inbred strains LSD from Pop1 and FD2M1 from admixed Pop1/Pop2

According to the breeding history and pedigree data, two strains were nearly 100% inbred: LSD (F > 0.99) and FD2M1 (0.98 < F > 0.99). However, based on the genetic data the residual heterozygosity in FD2M1 (0.33 SNP per kb on average) was still comparable with the less inbred strains ATHL and ATTHN (Table 1 and Figure 2A). This outcome provides a caution for the prediction of residual genetic variation levels based on inbreeding coefficients alone. In contrast, LSD individuals consistently have the lowest heterozygosity (0.00014), with very few segregating SNPs (Figure 2A), which is fully consistent with its calculated inbreeding coefficient of >0.99. We conclude that both LSD and FD2M1 (of which all living animals in August of 2021 have inbreeding coefficients of 0.996-0.997) are inbred strains, although FD2M1 had higher residual heterozygosity than LSD when the strains were sampled in 2014-2017, and might still have higher residual heterozygosity. A higher level of residual heterozygosity in the FD2M1 strain was expected, since the Pop1/Pop2 admixed FD2M1 strain certainly had a higher level of heterozygosity at the outset of the inbreeding program than the Pop1 LSD strain.

### Principal component analysis of the genetic data

To investigate population genetic relationships among the eight *M. domestica* strains, we performed principal component analysis (PCA) using the genotypes of 66,640 SNPs in the 70 ddRAD-seq individuals. The partially inbred sibling strains FD2M1 and FD2M4 clustered together, and they are distantly related to other strains on PC1, which is consistent with their admixed Pop1/Pop2 genetic background (Figure 3A). LSD animals are well separated from other strains by PC2, presumably through inbreeding and divergence from the LL1 random-bred strain (Figure 3A). The LL1-derived partially inbred sibling strains (AH11L, ATHHN, and ATHL) clustered together, with the more inbred ATHL strain completely separated from AH11L/ATHHN animals. This result is consistent with the fact that the AH11L and ATHHN strains were established as sibling strains well after the ATHL strain was established from the common founders of the three strains. LL1 animals are grouped in the center of the PCA plot, with the FD8X cluster next to them (Figure 3A). This is consistent with the fact that FD8X is a 1:1 mixture of Pop5 and LL1 background (Table 1). The same population genetic structure pattern remained evident in the PC plot after adding the whole-genome resequencing data from six additional FD8X and three additional LSD animals (Figure 3B).

**Figure 3.**
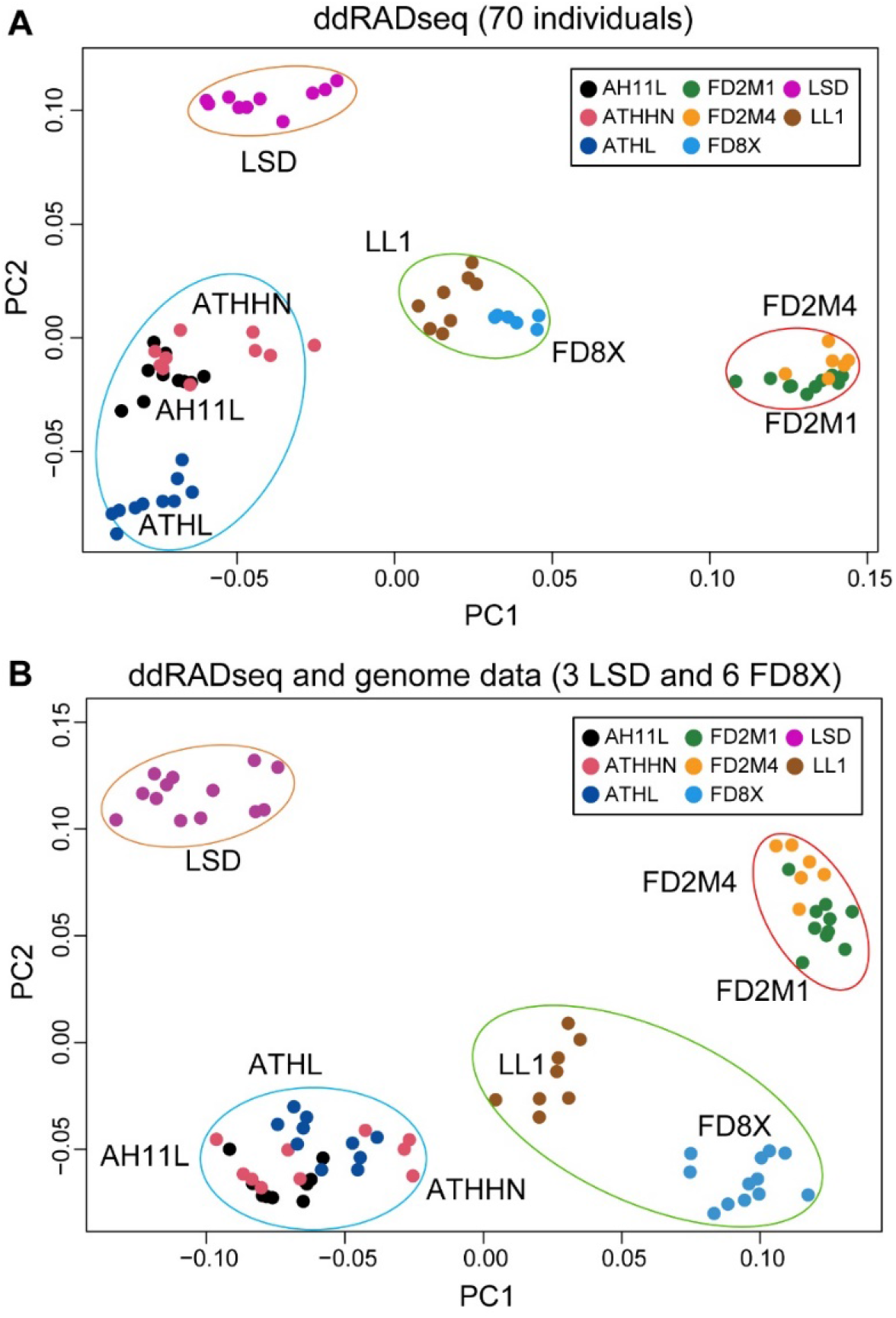
Principal component analysis (PCA) of genetic variations for the eight laboratory opossum strains. Scatterplot of individual variation in principal component (PC) scores (PC1 on the *x*-axis, PC2 on the *y*-axis) for 66,640 SNP loci of ddRAD-seq data in 70 individuals from eight strains **(A)** and the 70 ddRAD-seq data combined with 9 additional whole-genomic resequencing samples **(B)**.

### Population structure and admixture

In order to determine whether the population genetic structure agrees with the breeding and admixture history of these eight strains, phylogenetic, shared co-ancestry, and admixture analyses were performed using genotyping data from the 66,640 ddRAD SNPs. The phylogeny showed a deep split between the admixed Pop1/Pop2-derived FD2 group (FD2M1 and FD2M4) and LL1 and other Pop1 strains (LSD, AH11L, ATHHN, and ATHL) (Figure 4A). In the admixture analysis, models with K = 2, 3, or 4 were applied, and the eight strains were assigned to distinct groups at K=3 and K=4 (Figure 4B). FD2M1 and FD2M4 individuals are intermingled in the phylogeny (Figure 4A). They belong to the same population in the admixture analysis (Figure 4B), and the co-ancestry analysis revealed no differentiation between the two strains (Figure 4C). Therefore, we conclude that they were already quite highly inbred and had limited heterozygosity at the time they were separated as sibling strains. Similarly, the AH11L and ATHHN sibling strains lack clear separation in the phylogeny and shared co-ancestry analysis, and there is little to no differentiation between them. The sibling ATHL strain was assigned to a different population at K=4, but not at K=3 (Figure 4B). LSD is well separated from other Pop1-derived strains because of long-term inbreeding and more distant common ancestry with the AH11L, ATHHN, and ATHL strains.

**Figure 4.**
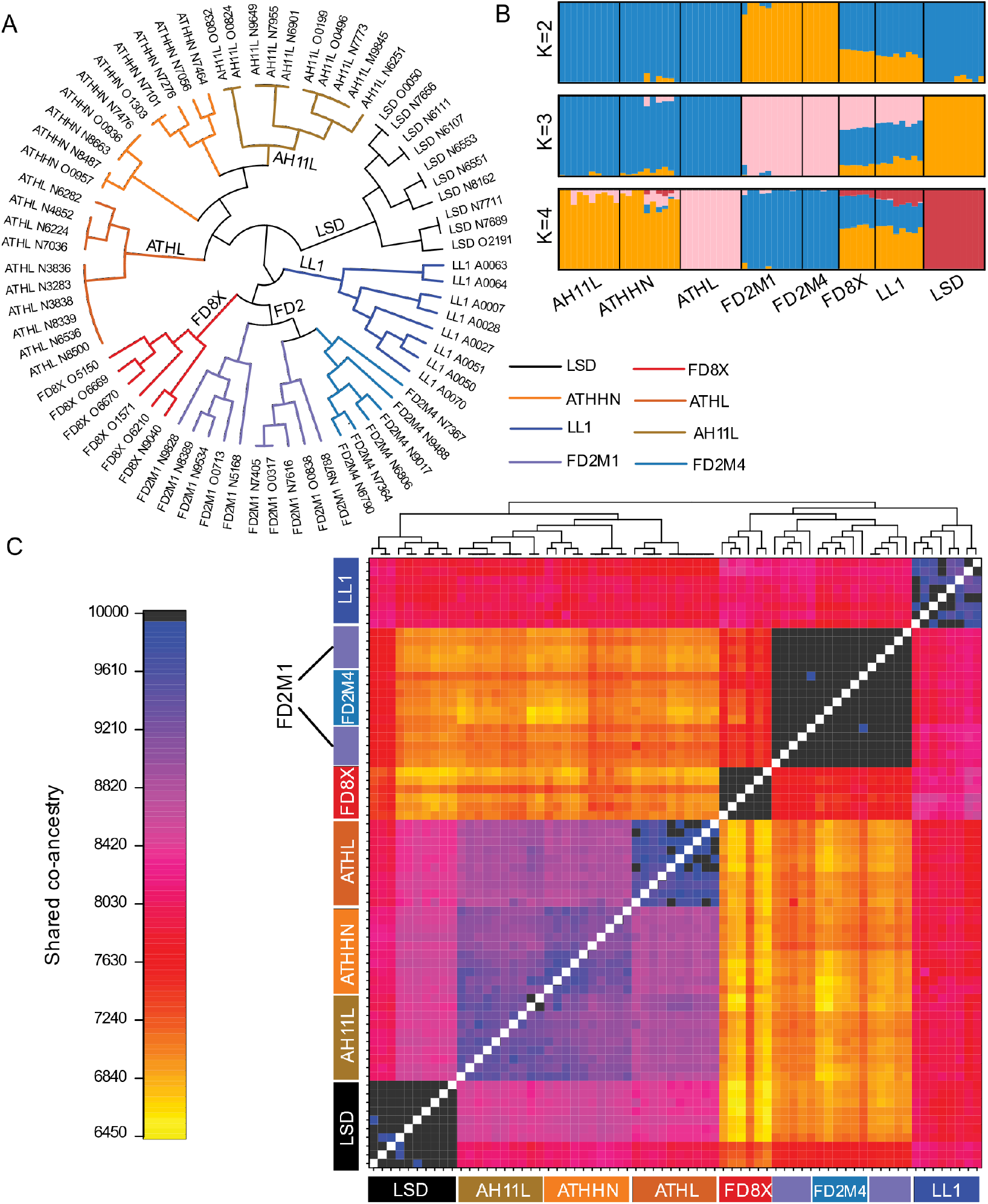
Graphical summary of genetic structure for ddRAD-seq individuals based on 66,640 SNP loci. **(A)** A phylogenetic tree was constructed using information from 66,640 SNPs. Tree edges represent individuals and are color coded as labeled. **(B)** Population genetic structure analysis plots with the number of subpopulations K = 2, 3, and 4. **(C)** The fineRADstructure plot showing population structure via shared co-ancestry. The cladogram depicts a clustering of individuals based on the pairwise matrix of co-ancestry. The heatmap depicts variation in pairwise co-ancestry among individuals according to the scale shown on the left. **(B)**

### Patterns of genetic diversity

We investigated the distribution of SNP density across all chromosomes and found that 1) it is elevated in telomeric relative to non-telomeric regions of the autosomes (Figure 5A), 2) the majority of the X chromosome has less than 1 SNP per kb (Figure 5A), and 3) there is a significant negative correlation between the inbreeding coefficient and the frequency of heterozygous SNPs (Spearman’s correlation coefficient ρ = −0.67, *P*-value = 2.5×10^−10^). Estimated nuclear diversity (π) is 0.0011 for the LSD strain and 0.0012 for the AH11L, ATHHN, and ATHL strains. The FD8X (F = 0.22) and LL1 (F = 0.42) strains were much less inbred than the other six strains examined (although some inbreeding has occurred due to the low number of founders), and had higher nuclear diversity (Figure 5C) and lower Fst values than the more inbred strains (Figure 5D). The genetic diversity patterns varied along the chromosomes, with elevated π in subtelomeric and subcentromeric regions (Figure 5C-D and S3-S4).

**Figure 5.**
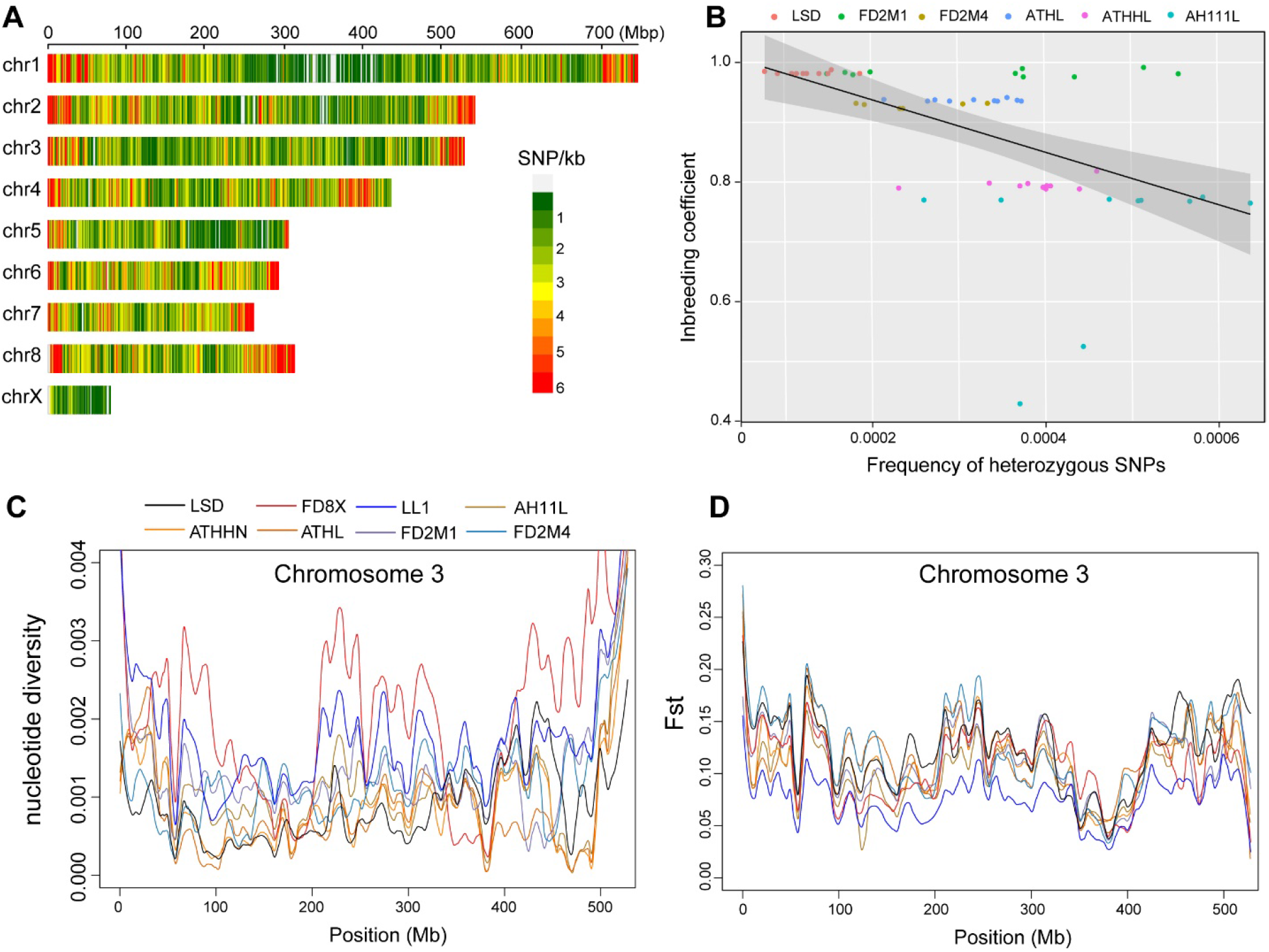
Patterns of genetic diversity across ddRAD-seq individuals from eight opossum strains. **(A)** SNP density plot showing the distribution of polymorphism density per million base pairs across chromosomes. The scale for the number of SNPs is shown on the right. **(B)** The correlation between inbreeding coefficients and frequencies of heterozygous SNPs in inbred strains. **(C)** Window-based nucleotide diversity along chromosome 3 for eight opossum strains (window size = 20kb). **(D)** Pattern of Fst value along chromosome 3. The strains are color-coded as described in the legend of Figure 5C.

## Discussion

Metatherians, or marsupial mammals, diverged from eutherian mammals ~160 million years ago and serve as vital models for research in comparative genomics and medicine. The gray short-tailed opossum (*M. domestica*) is the only well-established small laboratory marsupial model suitable for biomedical research, due to the advantages of non-seasonal breeding, relatively rapid sexual maturation, availability of multiple genetic stocks, and a high-quality reference genome. Although the genome sequence of one animal was published in 2007, the genetic diversity of different *M. domestica* laboratory strains has not previously been characterized. To fill this gap, our research provided a comprehensive analysis of the genetic architecture in eight laboratory strains.

The first generation *M. domestica* linkage map was constructed using the GMBX mapping panel, which consisted of F1 and F2 progeny of Pop1 and Pop3 crosses (Figure 1A) (Samollow et al. 2004). To improve the marker density and resolution, a second (Brazilian/Bolivian back-cross) mapping panel, BBBX, was established using crosses between Pop1 and Pop5 animals, which were descended from founders collected in Bolivia (Figure 1A) (Samollow et al. 2007). Pop5 is the most geographically distant of the five populations, and FD8X, which was established through 1:1 admixture of Pop5 and LL1, is the only laboratory strain with Pop5 ancestry (Figure 1B). We expected to discover significantly more SNPs in FD8X individuals compared to the admixed Pop1/Pop2 FD2M strains; however, we found SNP density in FD8X to be quite similar to those in the FD2M strains (Figure 2). There are several possible explanations, which are not mutually exclusive. First, the FD8X strain has experienced several near-loss events during its history, and these severe bottlenecks could be responsible for genome-wide SNP loss. Second, due to the large geographic distance between the Pop1 and Pop5 source populations, there could be incompatibilities between Pop1 and Pop5 genomes that lead to non-random losses of genomic segments from one or the other populations after the admixture. Some SNPs in tight linkage disequilibrium with such incompatible segments would also have suffered lost heterozygosity or fixation. Third, and considered by the authors to be least likely to be correct, FD8X SNPs could be underrepresented in the represented genomic regions just by chance, since the ddRAD-seq is a reduced representation sequencing approach.

The availability of the inbred strains, LSD and FD2M1, provides an essential genetic toolkit that enables strain-specific SNP differences to be used as positional tags for investigating a wide variety of allele-specific genome structures and gene expression characteristics. For example, documented genetic differences between these stains can be used to investigate allelic imbalances in gene expression DNA methylation, and histone modifications genome-wide as a means to better understand important phenomena of genomic imprinting, X-chromosome inactivation, and *cis*- vs. *trans*- regulation of gene expression (Wang and Clark 2014). As inbred strains are ideal for controlling genetic background in various kinds of experimental design, the past absence of inbred laboratory opossum strains hampered the potential research applications of this marsupial model. There have been multiple attempts to develop inbred strains of *M. domestica*, most of which have failed due to reduction in fertility and litter survival, presumably caused by inbreeding depression. In contrast, the inbred LSD strain is moderately fertile with an average litter size at the time of birth of 7, which compares favorably with a mean litter size of 9 for LL2, the most fertile of all stocks. In our ddRAD-seq data, we confirmed that the LSD animals consistently have the lowest residual heterozygosity compared to other inbred strains (1 per 10 kb in 2014). We are confident that LSD will prove to be an excellent resource for controlled treatment studies such as toxicological, developmental, and immunological research.

The FD2M1 strain, with a mean inbreeding coefficient of 0.985 at the time of sampling, was nearing the state of being fully inbred (F > 0.99). It, too, has continued to be further inbred, and all living FD2M1 animals currently (August, 2021) have inbreeding coefficients of 0.996-0.997. By comparison with LL2, this strain has a low level of fertility, with an average litter size of 5. Since its genetic makeup is quite distinct from the genetic makeup of the LSD strain, this pair of inbred strains can serve to determine the comparative effects of experimental treatments on two distinct genetic backgrounds, with minimal individual variation within each genetic background. Moreover, he two strains provide opportunities for rigorous genetic research on characters of biological and medical interest by conducting intercrosses and backcrosses, in which virtually all of the allelic differences between the two parental strains are known. Thus, our results have vastly increased the potential utility of this unique animal model.

Recently Kiyonari *et al.* (2021) reported breakthroughs in reproductive manipulation techniques for *M. domestica* that enabled them to achieve the first targeted gene perturbations through CRISPR/Cas9 genome editing in any marsupial species (Kiyonari et al. 2021). The ability to generate knockout animals, together with the availability of two inbred strains (with promise of more such strains as our inbreeding program continues), opens the way for investigating specific gene functions in *M. domestica* through knockouts of different alleles of a locus in two distinct and highly uniform genetic backgrounds.

In conclusion, our research is the first step in developing a strain-specific genomic toolkit for this marsupial model. We performed ddRAD-seq in eight *M. domestica* laboratory strains and investigated the genomic diversity within and between them. Population genetic parameters were computed based on 66,640 high-quality SNPs among these strains, and PCA and admixture analysis revealed that the population genetic structure is consistent with the original geographic locations of the founder animals and the breeding history since the introduction of the founders into the laboratory. We predict that the intersection of genetically diverse inbred strains and CRISPR/Cas9 gene-altering technologies will greatly enhance the utility of *M. domestica* for comparative biological and biomedical research, and will usher in a new era of scientific discovery based on this unique laboratory marsupial.

## Data availability

The whole-genome resequencing raw reads are available at NCBI SRA databases under accession number PRJNA743944. The ddRAD-seq raw data are available at NCBI SRA databases under accession number PRJNA743991. The SNP genotypes are available at https://github.com/XuWangLab/2021_ddRADseq_sppData.

## Acknowledgments

X.W. is supported by a USDA National Institute of Food and Agriculture Hatch project 1018100, National Science Foundation EPSCoR RII Track-4 Research Fellowship (OIA1928770), an Alabama Agriculture Experiment Station (AAES) Agriculture Research Enhancement, Exploration, and Development (AgR-SEED) award, and laboratory start-up and Research Support Program (RSP) funds from Auburn University College of Veterinary Medicine. X.X. and W.C. are supported by the Auburn University Presidential Graduate Research Fellowship and College of Veterinary Medicine Dean’s Fellowship. W.C. is supported by Alabama EPSCoR Graduate Research Scholars Program. We acknowledge the Auburn University Easley Cluster for support of this work. At TAMU, we thank Dr. William J. Murphy and Jonathan Bravo for DNA extraction and the Department of Veterinary Integrative Biosciences for animal support. At UTRGV, we thank Susan Mahaney for technical support and the Department of Human Genetics for support of the laboratory opossum research resource.

## Supplemental Materials

### Supplemental Figures

**Figure S1.** Histograms of proportions of homozygous/ heterozygous SNPs for eight autosomes of 70 individuals in eight opossum strains.

**Figure S2.** Barplot of numbers of heterozygous SNPs in female (left) and male (right) lab opossum samples.

**Figure S3.** Window-based nucleotide diversity plots along autosomes and the X chromosome for eight laboratory opossum strains (window size = 20kb).

**Figure S4.** Window-based Fst plots along autosomes and the X chromosome for eight laboratory opossum strains (window size = 20kb).

### Supplemental Tables

**Table S1.** Summary of raw read counts, quality control statistics, and genome mapping percentages for ddRAD-seq individuals.

**Table S2.** Summary of raw read counts, quality control statistics, and genome mapping percentages for the nine whole-genome resequencing individuals.

**Table S3.** Inbreeding coefficients calculated based on pedigree information for ddRAD-seq individuals.

**Table S4.** Population genetic parameters for the eight *Monodelphis domestica* strains.

### Supplemental Data

**Data S1.** *Monodelphis domestica* SNP ID, position, and genotype in ddRAD-seq and whole-genome resequencing individuals.

**Data S2.** SNP genotypes in RNA-seq data and ddRAD-seq data for 2,151 overlapped SNP positions.

